# Macroecological variation in movement profiles: body size does not explain it all

**DOI:** 10.1101/2022.04.21.489049

**Authors:** Samantha Straus, Coreen Forbes, Chelsea J. Little, Rachel M. Germain, Danielle A. Main, Mary I. O’Connor, Patrick L. Thompson, Adam T. Ford, Dominique Gravel, Laura Melissa Guzman

## Abstract

Animals couple habitats by three types of movement: dispersal, migration, and foraging, which dynamically link populations, communities, and ecosystems. Spatial distances of movement tend to correlate with each other, reflecting shared allometric scaling with body size, but may diverge due to biomechanical, phylogenetic, and ecological constraints. While these constraints have been investigated within specific taxa, the macroecological and macroevolutionary constraints on movement distances, and causes of those constraints, are still unknown. Here, we synthesized distances of all three movement types across 300+ vertebrate species, and investigated how the relationships between movement types and body size were modified by movement medium, taxonomy, and trophic guild (carnivore, herbivore, etc.). We found that the strength of relationships between movement types and body size varied among environments, taxa, and trophic guilds. Movement profiles interacted with physiological, taxonomic, and ecological traits to depart from expected body mass scaling. Overall, we find that there are systematic patterns to movement distances, and that movement types with very distinct ecological consequences (foraging, migration) can be correlated and subject to similar constraints. This implies that the scales of population dynamics in ecological communities are not entirely determined by the environment and likely reflect general biomechanical, evolutionary and metabolic constraints.

## Introduction

Movement within and between habitats is key to the persistence and stability of populations (Levins 1969), the diversity of communities (MacArthur & Wilson 1967; Leibold *et al*. 2004), the spatial distributions of species (Pulliam 2000), and the flow of energy, matter, and information between ‘donating’ and ‘receiving’ ecosystems (Loreau *et al*. 2003). Most organisms must move to forage, disperse to find mates, and migrate to thermoregulate, reach breeding grounds, track resources and escape predation. Each of these activities occur at distinct temporal and spatial scales (Box 1). Together, these movement types combine to create a multivariate ‘movement profile’ (a subset of ‘spatial use properties’ summarized by Guzman et al. (2019)) (Figure 1c-e). Though the importance of movement for biodiversity and the scales at which ecological systems change is becoming clear, it remains empirically very challenging to make a priori estimates of movement distances, and therefore to predict their consequences, for large groups of organisms. We contend that quantifying movement profiles across taxa will not only provide a deeper understanding of the inherent constraints to which animal movement are subject, but additionally, would focus a macroecological lens onto how and why movement types are distributed as they are among taxa (analogous to the plant trait spectrum in Diaz et al. (2016)). Quantified movement profiles would also provide a baseline to parameterize metacommunity and landscape models that link spatial dynamics to biodiversity patterns.

**Figure 1.**
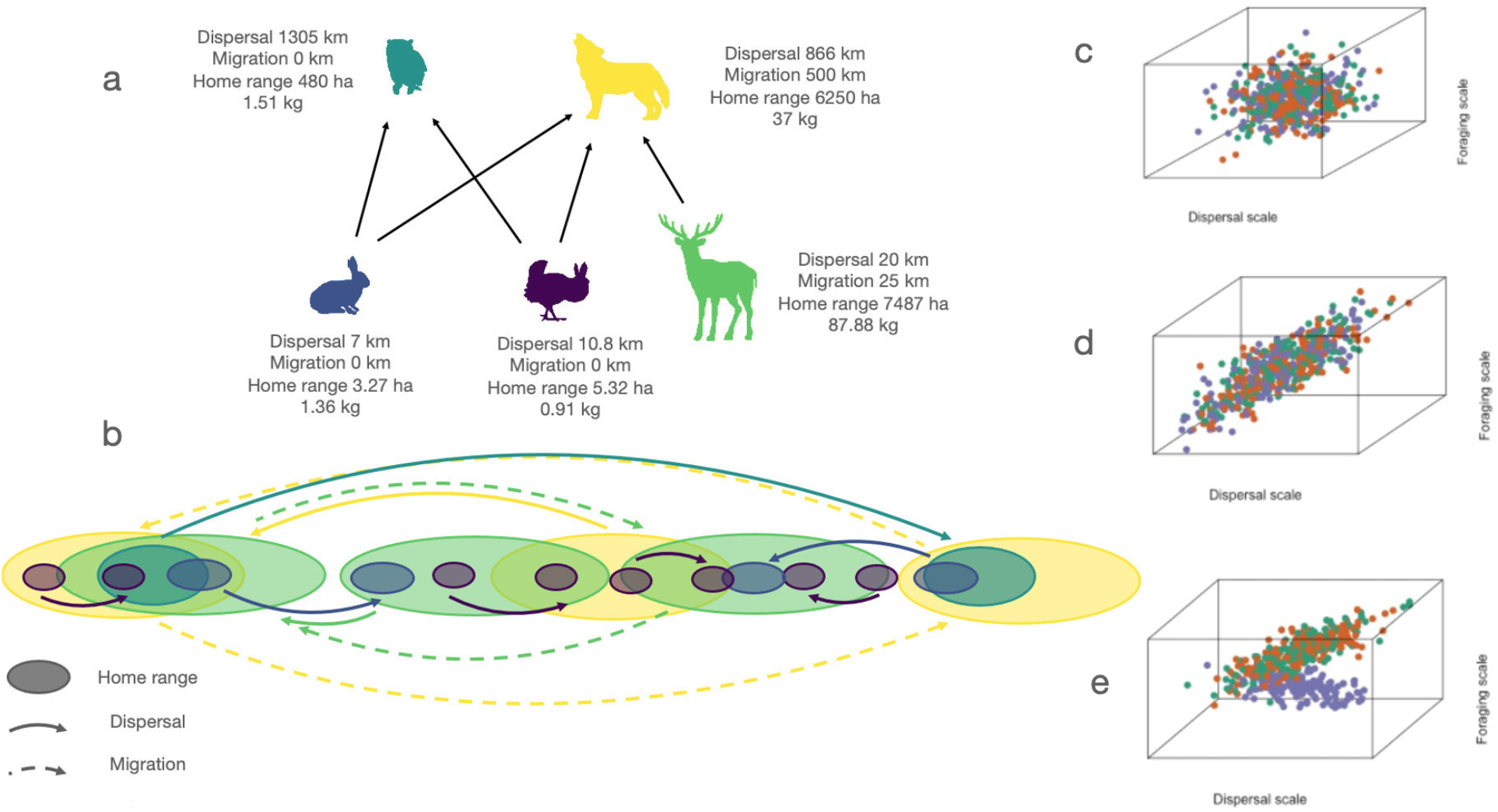
(a) A hypothetical food web with three herbivores and two carnivores. Each species has its own mean dispersal, migration, and foraging (home range) distance, and body size. (b) Each of the three types of movements influence when and where members of the food web interact. Points in plots c-e represent the hypothetical trivariate movement profiles of a species, and color representing a constraint with three levels. (c) Under the null hypothesis, movement types do not covary. (d) three movement types covary strongly because movement scales with body size. (e) certain groups, either due to evolutionary or ecological constraints, diverge from expected patterns of covariance between movement types

While movement profiles are expected to scale with body mass, deviations from the expected scaling can reflect ecological processes that constraint movement. Body size scales allometrically with metabolic demands, and consequently, mass-specific costs of movement decline with increasing body size (Schmidt-Nielsen 1972; Shepard *et al*. 2013). It follows that larger animals may be able to move farther per unit mass than smaller ones, setting the spatial scale and resolution of population dynamics at a coarser scale as body size increases. Several studies report that, for each movement type individually, movement covaries with body size (Hein *et al*. 2012; Santini *et al*. 2013; Tamburello *et al*. 2015). We considered, as a null hypothesis, that characteristic distances associated with all three movement types covary with body size, and thus, each other, thereby constraining possible scales of movement and its dynamic consequences (Table 1, Figure 1). However, movement scaling may depart from expected body mass scaling due to biomechanical, phylogenetic, and ecological constraints. Understanding these additional constraints on movement is key to accurately predicting the role movement processes play in ecology.

**Table 1.** Table with listed hypotheses

| Hypothesis | Description | Prediction |
| --- | --- | --- |
| **H0: Body size** | Covariance between movement types results from allometric scaling | Positive covariance between movement types and body size |
| **H1: Movement medium hypothesis** | Some media (e.g. water) will be harder to move through than others (e.g. air) | Different intercepts (movement type ~ body size) for different media, decoupling of covariance (e.g. different substrate penetrability of different media) |
| **H2: Shared taxonomy hypothesis** | Covariance results from shared taxonomic history | Expect taxonomy to decouple covariance between movement types, e.g. phylogenetic inertia |
| **H3: Trophic guild hypothesis** | Foraging movement should be linked with trophic guild, as carnivores may forage over larger distances. Other movement types may not be affected by trophic guild | Expect diet to decouple covariance between foraging and other movement types, e.g. higher trophic levels moving farther |

Our study considered three constraints on movement: movement medium, shared taxonomy, and trophic guild. First, as a biomechanical constraint, movements through different media (i.e. air, water, land) incur different energetic costs, reflecting physical factors such as viscosity, drag and gravity (“substrate penetrability”, (Shepard *et al*. 2013)). Movement media can influence the energy landscape an animal encounters, where inclines, drafts, and currents can all relieve or exacerbate the cost of transportation (Shepard *et al*. 2013; Gallagher *et al*. 2017). Second, our null hypothesis is that movement profiles are all causally dependent on body size. In this scenario, as body sizes have evolved over macroevolutionary time (whether gradually or in large jumps in trait space (Landis & Schraiber 2017)), we expect the movement profiles will evolve in response with little evolutionary inertia (Hansen & Orzack 2005); as such, once we condition on body size, there should be no residual phylogenetic signal in movement profiles (Uyeda *et al*. 2018). However, if there are other unmeasured variables that influence movement profiles (i.e., aside from body size) and if these have phylogenetic signal themselves, we may see variation in the scaling relationship between body sizes and movement profiles among clades (Uyeda *et al*. 2017). This could lead to departures from the expected scaling of movement with body mass. Finally, movement may be constrained by the ecological roles of species within food webs (e.g., their trophic guild, or whether they are carnivores, herbivores, insectivores or omnivores). Higher trophic levels may move across longer distances to meet energy requirements (Harestad & Bunnel 1979), different trophic guilds may exhibit unique movement profiles. Empirical estimates of these relative spatial extents of movement are required to extend spatial diversity theories (e.g., metacommunity ecology, spatial coexistence theory) into a multi-trophic context, where interactions (e.g., predation, mutualism) are often realized between species with vastly different body sizes, ecologies, and evolutionary histories (Guzman *et al*. 2019).

We synthesized movement data from databases and published studies for 322 species of animal to test the following hypotheses: (i) the ‘movement medium hypothesis’: movement through less penetrable media, e.g. water, results in shorter distances than movement through easily penetrable media, e.g. air; (ii) the ‘shared taxonomy hypothesis’: animals with shared taxonomy are similarly constrained in their movement profiles, and the covariance between movement types and body mass change based on the taxa considered, i.e. signals of “unreplicated evolution events”; and (iii) the ‘trophic guild hypothesis’: trophic guild, a trait of fundamental importance in ecological systems, influences movement such that carnivores move longer distances than other guilds, e.g. herbivores (Table 1). If our null hypothesis that movement distance scales with body size is supported, the covariance between movement types should disappear when we account for body size. Deviations from this expectation may arise when ecological (e.g. trophic level, medium of movement) or evolutionary forces interact to decouple the characteristic distances associated with each type of movement (Figure 1).

### Box 1.

***Movement types uniquely affect ecological dynamics***

**Foraging** refers to movement to acquire resources within a species’ home range, structuring trophic interactions. Larger animals and those at higher trophic levels may forage over larger areas to meet resource requirements, and in doing so, connect multiple habitats occupied by the organisms that they consume (McCann *et al*. 2005; McCauley *et al*. 2012). Of the movement types considered, foraging generally occurs at the shortest distances (Guzman *et al*. 2019). Likewise, foraging movements occur most frequently, up to multiple times per day for some organisms. Resource quality within a habitat patch affects the distance and duration of movement (Charnov 1976), where organisms foraging within low quality or high variability habitats move more often than those in high quality or low variability ones. Similarly, generalist species may not need to move as often or far as specialists. Competition within a trophic level may also be influenced by foraging movements. For example, movement into or out of an area may induce or release the negative effects of one population on another’s growth rate via competition over shared resources or enemies (Amarasekare 2008). Here we use home range size as a proxy metric for foraging movement, because it encompasses foraging movements and is most consistently measured for multiple taxa.

**Dispersal** is the once-in-a-lifetime movement of individuals between habitat patches. Interspecific variation in dispersal distance is generally greater than for the other forms of movement (e.g. blue sharks >7500 km vs. common musk turtle <100m (Jenkins *et al*. 2007)). Dispersal distances within species often display strongly right skewed distributions, such that most individuals disperse short distances and a few disperse very far (Nathan *et al*. 2003). Dispersal may be widespread among individuals within the natal stage followed by a more sessile adult stage, may be sex biased, or may occur at any life stage as a response to population density or suboptimal conditions. Dispersing individuals can colonize habitat patches that were previously unoccupied, increasing the geographic range of that species (Holt & Keitt 2005), or they can move to different areas of an occupied range, thereby demographically and genetically connecting populations. Because of this role in population dynamics, dispersal is the most commonly considered movement type in population and community ecology (Hubbell 2001; Vellend 2010). Importantly, “immigration” and “emigration”, when used in the context of population genetics and community ecology, usually refer to dispersal away from or into a habitat (as opposed to actual ‘migration’).

**Migration** refers to cyclical (e.g. seasonal) movements to track resources or mates, and thus has a regular and unique temporal structure. Migration ranges from hundreds of meters (for amphibians accessing breeding ponds) to hundreds of kilometers (ungulates tracking food). Migration occurs one or more times in an organism’s lifetime, depending on a species’ life history. Rather than connecting different populations, it connects different habitats used by the same population, as well as places along a migratory route. Migration is a necessary strategy for persistence (Sinclair 2003), but remains understudied in metapopulation and metacommunity ecology, despite its trophic and competitive effects on donor, recipient, and en-route communities through which animals move (Cohen & Satterfield 2020). Instead, migration is of particular interest to ecosystem and meta-ecosystem ecology, as migratory species can transport nutrients over vast distances, affecting the functioning of far-flung ecosystems (Bauer & Hoye 2014; Gounand *et al*. 2018). In addition, migratory movements are often undertaken by entire populations or aggregations of animals, which can engineer the timing and magnitude of ecosystem dynamics such as primary production (Geremia *et al*. 2019) or create large, temporally discrete pulses of nutrients (Subalusky *et al*. 2017).

## Methods

### Data sources

To test our hypotheses about whether distances of different movement types (dispersal, migration, foraging) are related and whether they reflect biomechanical, ecological or other constraints, we synthesized observations on each of the three movement modes for taxa comprising major vertebrate taxonomic groups, a wide range of body sizes, and representing aquatic and terrestrial life histories. We drew upon existing databases and individual empirical studies (see Supplementary References, aiming to assemble a taxonomically diverse dataset. We drew upon home range databases for mammals and birds (Armstrong 1965; Tamburello *et al*. 2015) and amphibians, migration databases for all taxa (Hein *et al*. 2012; Trochet *et al*. 2014), and dispersal databases for reptiles and amphibians (Jenkins *et al*. 2007; Trochet *et al*. 2014) and mammals (Santini *et al*. 2013). For species to be included in our dataset, an estimate for each of the three movement types had to be available, as well as data for body mass.

Once all databases were exhausted, we conducted a systematic search using Web of Science Core Collection (licensed to University of British Columbia) that filtered by English language, article document types, and citation indices from 1900 to present. We used the following search terms: (Set 1: Topic = (migration OR dispersal OR home range); Set 2: Topic = (meta analysis OR database); Set 3: Set 1 AND Set 2). Our search included the following categories: behavioral sciences, biodiversity conservation, biology, ecology, entomology, evolutionary biology, fisheries, limnology, marine freshwater biology, ornithology, zoology. If individual values were found from both existing published databases and from our Web of Science search, we chose the former. Finally, to fill in individual missing values (e.g., the dispersal range of a species for which we had already found values for home range size and migration distance) after Web of Science results were exhausted, we used Google Scholar, IUCN, Encyclopedia of Life (EOL), Animal Diversity Web (ADW), and included government reports and theses alongside peer-reviewed articles. Each search was confined to the top five returned pages.

### Data standardization

Several decisions were made during data collection to ensure high data standards and comparability across taxa, for each movement type. Dispersal distance was collected from experimental studies, observational studies, and meta-analyses. In order to relate movement to adult body size, adult median or mean dispersal distances (units: kilometers) were prioritized over juvenile, propagule, and natal distances. When median or mean distances were not provided, maximum dispersal distance was used (Nathan *et al*. 2003). When multiple means were provided within any one study, we calculated an average distance weighted by sample size. Active dispersal distances were chosen over passive dispersal, e.g., in the case of larval fish or amphibians. In rare cases (n=9), movement was estimated from maps (i.e. map of distance between two populations with gene flow to estimate dispersal). Migration distance (units: kilometers) was collected from experimental and observational studies. When more than one migration distance was provided by the same source, the largest distance was used. The weighted average was also used when migration distances were different between multiple individuals, or between males and females. For many non-migratory animals, we assumed a migration distance of 0 km. Likewise, foraging distance, estimated as the diameter of the home range area (assuming a circular home range, units: kilometers), was also collected from experimental and observational studies. Similar to migration, the largest foraging distance listed was used if more than one was given for a species, as well as the average between males and females.

We also collected data on predictors of movement that aligned with our hypotheses. To test our null hypothesis, body weight measured in mass (kg) was collected for all taxa (Armstrong 1965; Harestad & Bunnel 1979; Trochet *et al*. 2014; Tamburello *et al*. 2015);. When multiple body weights were provided, the average was used. Similarly, the average was taken for male and female measurements. To test the movement medium hypothesis, we characterized movement media for each movement type as land, water (aquatic or marine) or air or any combination of these media. Movement media were inferred for species with unambiguous locomotion. For example, the medium a hummingbird uses to locomote is air and for a shark it is water (marine). For most species, locomotion media were confirmed with a literature search. If more than one media significantly contributes to the movement of an animal, a combination of media were recorded. For example, an amphibian that reproduces in the water but lives on land might have land for medium of dispersal and migration and “land_aquatic” for their foraging medium. If a species engages in a form of locomotion only over extremely short distances compared to another form of movement, that movement media was not considered. For example, a grouse that forages on the ground and only flies in short bursts to avoid predators would have land as its medium. To test the ecological hypothesis, the trophic guild for each species was classified as herbivore, omnivore, invertivore, or carnivore. Data were gathered from the data papers and meta-analyses the movement data had been extracted from, where possible, and otherwise from the Encyclopedia of Life (https://eol.org), Animal Diversity Web (https://animaldiversity.org), or primary literature searches. To test the shared taxonomy hypothesis, we obtained Class and Order for each species using the package *taxize* using NCBI and ITIS databases (Chamberlain & Szöcs 2013).

### Statistical analyses

To explore correlations among movement distances among different movement types, and whether these correlations were related to movement medium, shared taxonomy, or trophic guild, we conducted principal component analyses (PCAs) for the tri-dimensional movement profiles of species using the *vegan* (2.5.7) (Oksanen *et al*. 2020) and *stats* (4.0.5) packages in R (4.0.5) (R Core Team 2021). For each proposed constraint, we conducted PCAs on both (i) movement distances (standardized and log10-transformed) and (ii) standardized residuals from a regression of movement distance as a function of log10 transformed body size. This second PCA was included to test our null hypothesis that movement types will be correlated due to underlying body mass scaling relationships.

Next, to estimate how each movement type was associated with body size and whether the relationship between movement type and body size was modified by other constraints, we used Bayesian generalized linear multivariate regressions using Stan through the *brms* package (Bürkner 2018). We chose this modelling approach because it allowed us to fit multivariate regressions to estimates of three response variables (dispersal, migration and foraging distance) simultaneously, while allowing different error distributions of each response variable. We used a Gamma response distribution with a log link function for dispersal and foraging distance, as these variables were continuously distributed while being bounded at zero. We used a hurdle gamma response distribution for migration because it allowed us to model non-migratory species (value of 0), as well as the distribution of the migratory species (values above of 0). These multivariate models had all three types of movements as a function of body size (Model 0 -null hypothesis).

For each hypothesized constraint (Table 1), we used random effects, allowing the relationships between body size and movement to vary among levels of each predictor variable (e.g. movement media types). The null model (Model 0) was compared to models including movement medium (Models 1.1-1.3), taxonomy (Model 2 using class for Model 2.1 and order for 2.2), and trophic position (Model 3) with intercept and slope random effects. By using random effects, each category within a hypothesis received a slope and intercept, but all of the slopes or intercepts across a model were all drawn from the same distribution. This type of model recognizes that the relationship between body size and movement is not completely independent for each category within a hypothesis. For media (Model 1), we conducted univariate analyses for each movement type in the media classification to account for species that use different media for different movements, e.g. waterfowl that forage aquatically but migrate aerially, and we excluded species from this analysis that used multiple media for one form of movement (Model 1.1, 1,2, 1.3). Models 2.1 and 2.2 test the effects of taxonomy through class (2.1) or order (2.2). For the main figures and tables we present the results of Model 2.1 (class), and we present results for model 2.2 in the supplementary materials. We natural log-transformed body mass for all models. We fit the following models:

Model 0 tests the effects of body mass. The same error structure was also used for models 1-3:

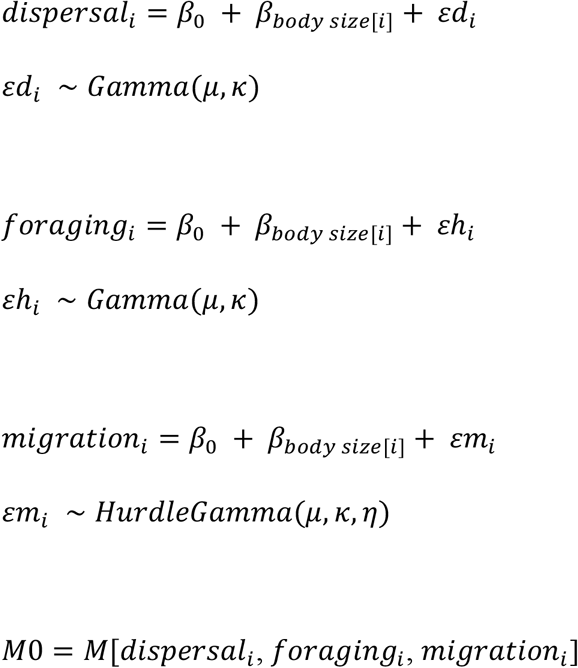

Models 1-3 followed a similar formulation, with random slope and intercept terms added for each constraint. E.g.:

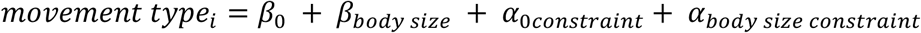

Models 2 and 3 were multivariate:

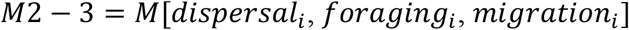

Model 1 was univariate because the movement media changed between movement types.

Full model formulations can be found in the supplemental materials. All of the models used the default uninformative priors, and they were run using 4 chains, using defaults for warm-up with no thinning. In order to improve model convergence, the acceptance rate (adapt delta) was increased to 0.995 and the maximum tree depth of the NUTS sampler to 15. Model convergence was assessed through visual inspection and where 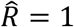. M0 used the default of 2,000 iterations; M1 used 5,000 iterations; M2.1 used 10,000 iterations; M2.2 used 5,000 iterations; and M3 used 3,000 iterations. To assess the performance Models 1 and 3 compared to the null model, we used the LOOIC information criterion. As Model 2 is composed of three univariate models, we compared slopes and intercepts to that of the null model. All data and code will be deposited in dryad upon acceptance.

## Results

We found movement distances for 322 taxa, a total of 966 observations. Birds and mammals were over-represented in the dataset relative to other taxa, with 159 and 116 species respectively, and the remaining 47 species were amphibians, reptiles, and sharks. Additionally, while our dataset included a few gliding mammals (e.g. flying squirrels), it lacked any volant mammals (e.g. bats and flying foxes) or insects, meaning there was complete overlap between animals using “air” as their medium of movement and the class Aves. Finally, while dispersal and foraging distances were unimodal, migration distances were strongly bimodal, with 138 species being entirely non-migratory (Figure 2). Initial examination of each potential constraint in the absence of the others indicated that species dispersed and foraged over greater scales in water than on land but migrated further through air (Figure 2a-c); that different taxonomic groups had different scales of movement (Figure 2d-f); and that unlike for media or taxonomy, species belonging to different trophic guilds had substantial overlap in their scales of movement (Figure 2g-i).

**Figure 2.**
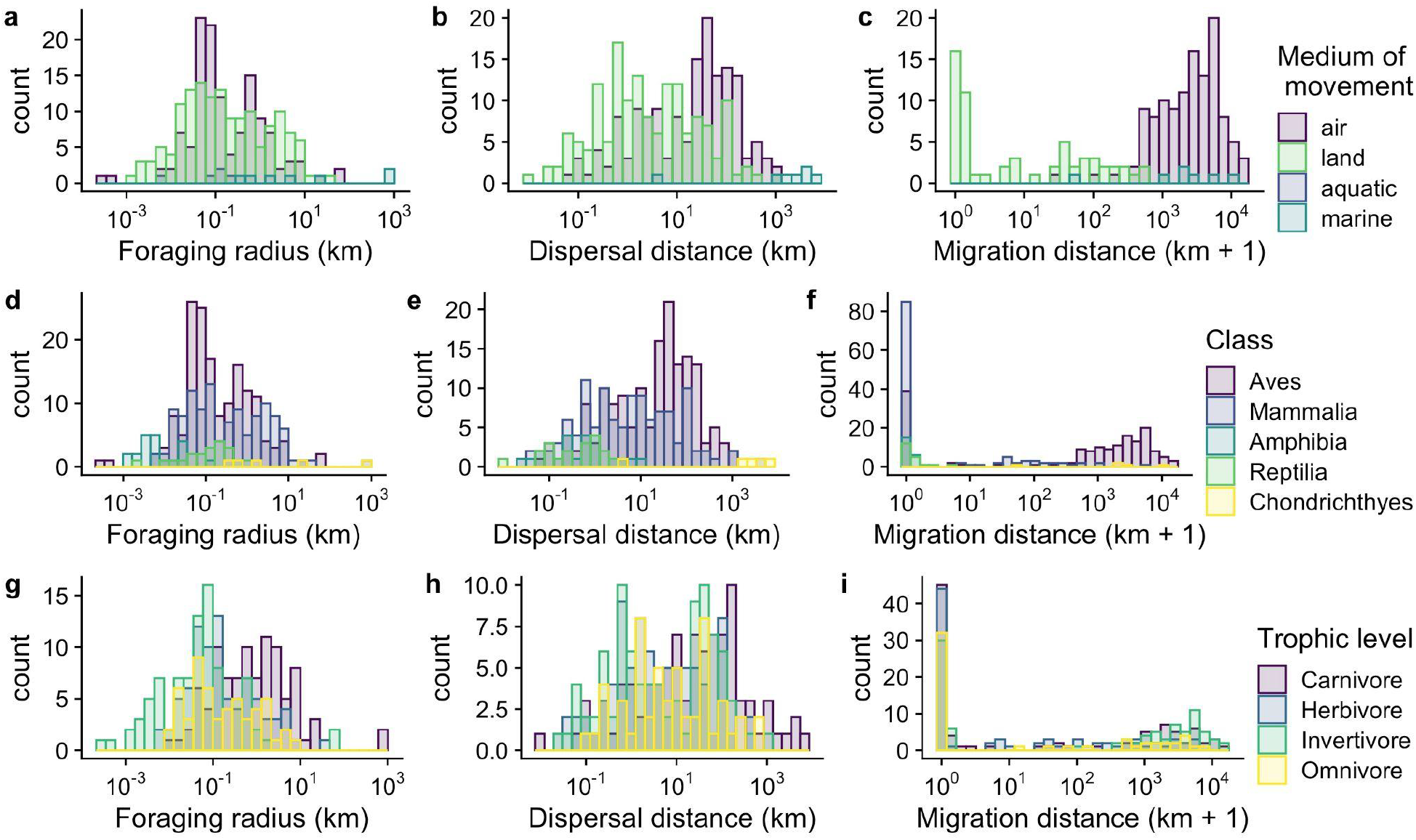
Distribution of SUPs by media (a-c), taxonomy (d-f), and trophic level (g-i). X-axes log10 transformed, migration log10+1

### Covariance between movement types

We found partial support for our null hypothesis that body size is, in part, responsible for covariation between movement types, however, body size was not the only source of this covariance. To test this null hypothesis, we draw upon two pieces of evidence. First, as predicted, foraging and dispersal distances were both positively correlated with body size (Table 2, Model 0), however in contrast to our predictions, migration distance was not. As an extension, we might expect controlling for body size might cause foraging and dispersal distances to become decoupled if body size were the main cause of their covariance. Indeed, considering all species together, this does appear to be the case: dispersal and foraging distances covaried prior to correcting for body size (i.e., similar loadings on PCA axes 1 and 2; Fig. 4) but less so afterwards (i.e., dissimilar loadings on PCA axes 1 and 2). This difference occurred despite the inclusion of body size having no effect on the total amount of variance in the dataset summarized by the first two PCA axes (85% with body size, 82% without body size). Interestingly, correcting for body size increased correlations between dispersal and migration distances (if considering only migratory species).

**Table 2.**
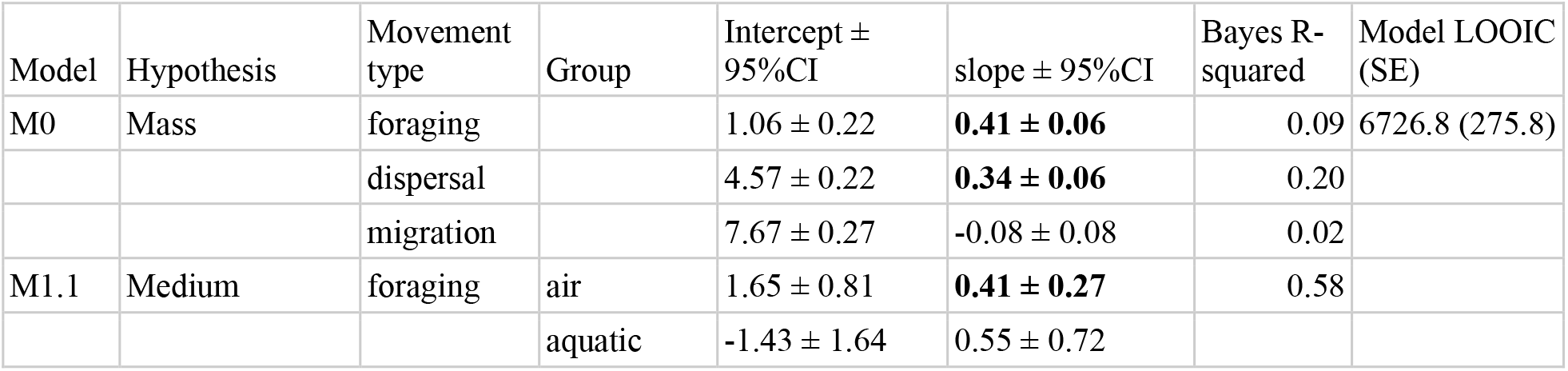
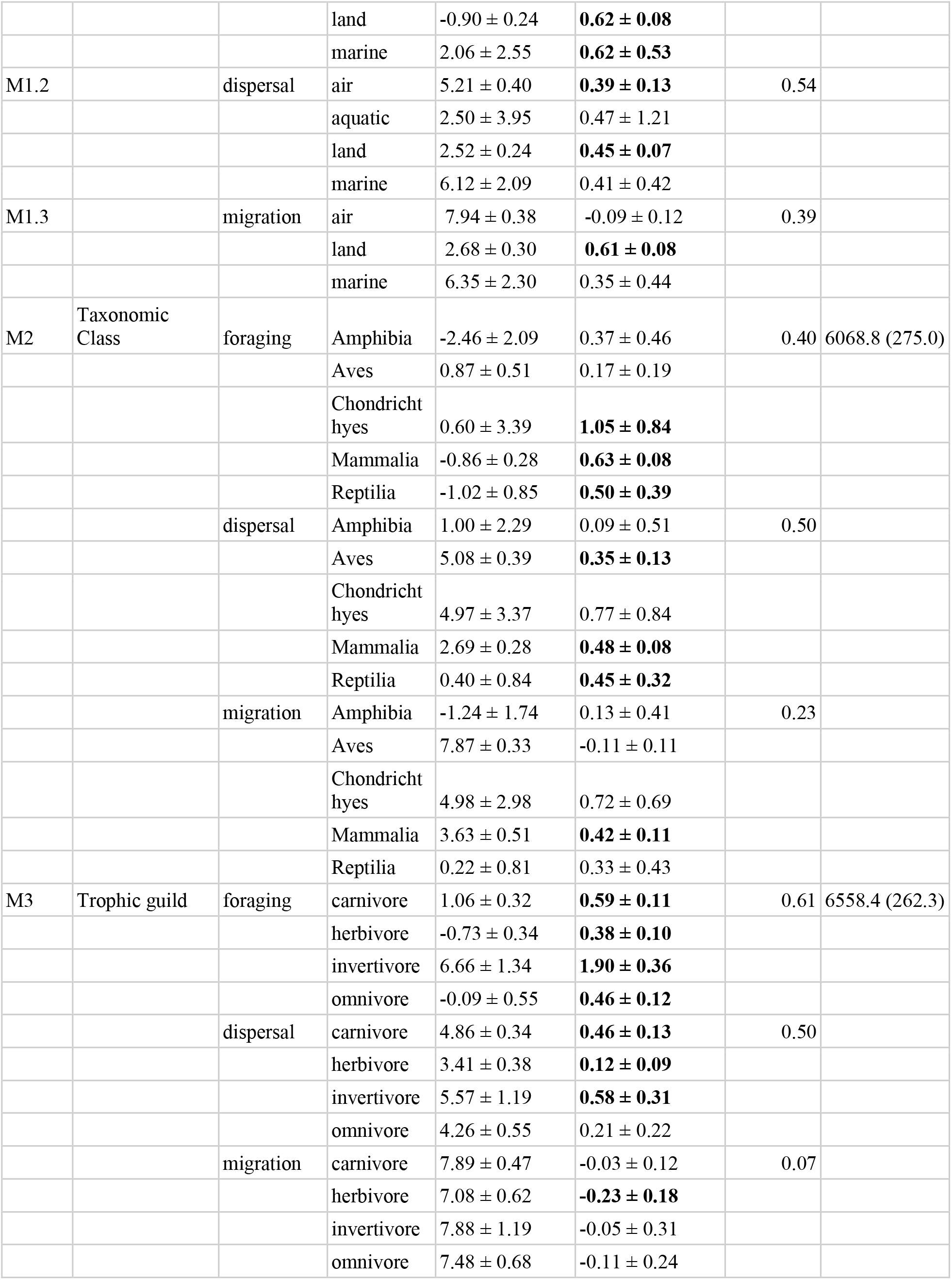
Slopes and intercepts for BRMS models including; mass, mass+class, mass+medium, mass+trophic level. **Bolded slopes** are significantly different than zero. For medium, each movement type was assigned to media separately to include species that may use different media for different purposes, e.g. foraging in water but migrating by air.

| Model | Hypothesis | Movement type | Group | Intercept ± 95%CI | slope ± 95%CI | Bayes R-squared | Model LOOIC (SE) |
| --- | --- | --- | --- | --- | --- | --- | --- |
| M0 | Mass | foraging | | 1.06 ± 0.22 | **0.41 ± 0.06** | 0.09 | 6726.8 (275.8) |
| | | dispersal | | 4.57 ± 0.22 | **0.34 ± 0.06** | 0.20 | |
| | | migration | | 7.67 ± 0.27 | -0.08 ± 0.08 | 0.02 | |
| M1.1 | Medium | foraging | air | 1.65 ± 0.81 | **0.41 ± 0.27** | 0.58 | |
| | | | aquatic | -1.43 ± 1.64 | 0.55 ± 0.72 | | |
| | | | land | $-0.90 \pm 0.24$ | **$0.62 \pm 0.08$** | | |
| | | | marine | $2.06 \pm 2.55$ | **$0.62 \pm 0.53$** | | |
| M1.2 | | dispersal | air | $5.21 \pm 0.40$ | **$0.39 \pm 0.13$** | 0.54 | |
| | | | aquatic | $2.50 \pm 3.95$ | $0.47 \pm 1.21$ | | |
| | | | land | $2.52 \pm 0.24$ | **$0.45 \pm 0.07$** | | |
| | | | marine | $6.12 \pm 2.09$ | $0.41 \pm 0.42$ | | |
| M1.3 | | migration | air | $7.94 \pm 0.38$ | $-0.09 \pm 0.12$ | 0.39 | |
| | | | land | $2.68 \pm 0.30$ | **$0.61 \pm 0.08$** | | |
| | | | marine | $6.35 \pm 2.30$ | $0.35 \pm 0.44$ | | |
| M2 | Taxonomic Class | foraging | Amphibia | $-2.46 \pm 2.09$ | $0.37 \pm 0.46$ | 0.40 | 6068.8 (275.0) |
| | | | Aves | $0.87 \pm 0.51$ | $0.17 \pm 0.19$ | | |
| | | | Chondrichthyes | $0.60 \pm 3.39$ | **$1.05 \pm 0.84$** | | |
| | | | Mammalia | $-0.86 \pm 0.28$ | **$0.63 \pm 0.08$** | | |
| | | | Reptilia | $-1.02 \pm 0.85$ | **$0.50 \pm 0.39$** | | |
| | | dispersal | Amphibia | $1.00 \pm 2.29$ | $0.09 \pm 0.51$ | 0.50 | |
| | | | Aves | $5.08 \pm 0.39$ | **$0.35 \pm 0.13$** | | |
| | | | Chondrichthyes | $4.97 \pm 3.37$ | $0.77 \pm 0.84$ | | |
| | | | Mammalia | $2.69 \pm 0.28$ | **$0.48 \pm 0.08$** | | |
| | | | Reptilia | $0.40 \pm 0.84$ | **$0.45 \pm 0.32$** | | |
| | | migration | Amphibia | $-1.24 \pm 1.74$ | $0.13 \pm 0.41$ | 0.23 | |
| | | | Aves | $7.87 \pm 0.33$ | $-0.11 \pm 0.11$ | | |
| | | | Chondrichthyes | $4.98 \pm 2.98$ | $0.72 \pm 0.69$ | | |
| | | | Mammalia | $3.63 \pm 0.51$ | **$0.42 \pm 0.11$** | | |
| | | | Reptilia | $0.22 \pm 0.81$ | $0.33 \pm 0.43$ | | |
| M3 | Trophic guild | foraging | carnivore | $1.06 \pm 0.32$ | **$0.59 \pm 0.11$** | 0.61 | 6558.4 (262.3) |
| | | | herbivore | $-0.73 \pm 0.34$ | **$0.38 \pm 0.10$** | | |
| | | | invertivore | $6.66 \pm 1.34$ | **$1.90 \pm 0.36$** | | |
| | | | omnivore | $-0.09 \pm 0.55$ | **$0.46 \pm 0.12$** | | |
| | | dispersal | carnivore | $4.86 \pm 0.34$ | **$0.46 \pm 0.13$** | 0.50 | |
| | | | herbivore | $3.41 \pm 0.38$ | **$0.12 \pm 0.09$** | | |
| | | | invertivore | $5.57 \pm 1.19$ | **$0.58 \pm 0.31$** | | |
| | | | omnivore | $4.26 \pm 0.55$ | $0.21 \pm 0.22$ | | |
| | | migration | carnivore | $7.89 \pm 0.47$ | $-0.03 \pm 0.12$ | 0.07 | |
| | | | herbivore | $7.08 \pm 0.62$ | **$-0.23 \pm 0.18$** | | |
| | | | invertivore | $7.88 \pm 1.19$ | $-0.05 \pm 0.31$ | | |
| | | | omnivore | $7.48 \pm 0.68$ | $-0.11 \pm 0.24$ | | |

By grouping species according to their movement media, taxonomy, and trophic guild, we can evaluate alternative hypotheses for causes of covariance between movement types unrelated to body size. After correcting for body size, we found the strongest support for the ‘biomechanical’ and ‘shared taxonomy’ hypotheses, but no support for the ‘trophic guild’ hypothesis, as causes for covariance among movement types unrelated to body size. In all cases, separation happened primarily along PCA axis 1, as different groups generally represented species that moved more or moved less overall, as opposed to moving more or less for specific movement types. Nonetheless, one of our most striking results is the fact that controlling for body size either reduced (e.g., movement media, taxonomic) or completely collapsed (e.g., trophic level) differences among groups that were otherwise pronounced on both PCA axes.

### Covariance and movement profile in three-dimensional trait space

By viewing all three dimensions of species’ movement profiles at once, we can better visualize the covariance structure described earlier, but additionally, pull out additional features that inform us about constraints on movement. First, two major areas of the volume were unfilled by any species in our dataset: areas characterized by long foraging distances and short dispersal distances, and the converse, short foraging distances with long dispersal distances. Second, movement profiles for several species had extreme values relative to the rest of the dataset, most notably, four species: two with extreme distances for all movement types (i.e., *Physeter catodon* (Sperm whales) - Mammalia, *Prionace glauca* (Blue shark) - Chondrichthyes) and two with exceptionally long (i.e., *Rangifer tarandus* (Reindeer) *-* Mammalia) and short (i.e., *Asio flammeus* (Short-eared owl) - Aves) foraging distance for a given dispersal distance. Third, given that different movement types were quantified in comparable km units, foraging radius had the widest range of values, spanning approximately 12 orders of magnitude, greater than dispersal (6 orders) and migration (5 orders) owing equally to extremely high (10^6^ km) and extremely low (10^−6^ km) foraging distances. Importantly, movement distances in Figure 3 are shown in log-space, meaning that the data shown would be skewed towards high values on a raw scale. When considering the boundaries of our three-dimensional trait space, we found that foraging and dispersal distances had uneven boundaries, particularly for migrating species. In contrast, boundaries on migration distances were more defined, both at the lower bound (for obvious reasons, as several species do not migrate) and also at the upper bound.

**Figure 3.**
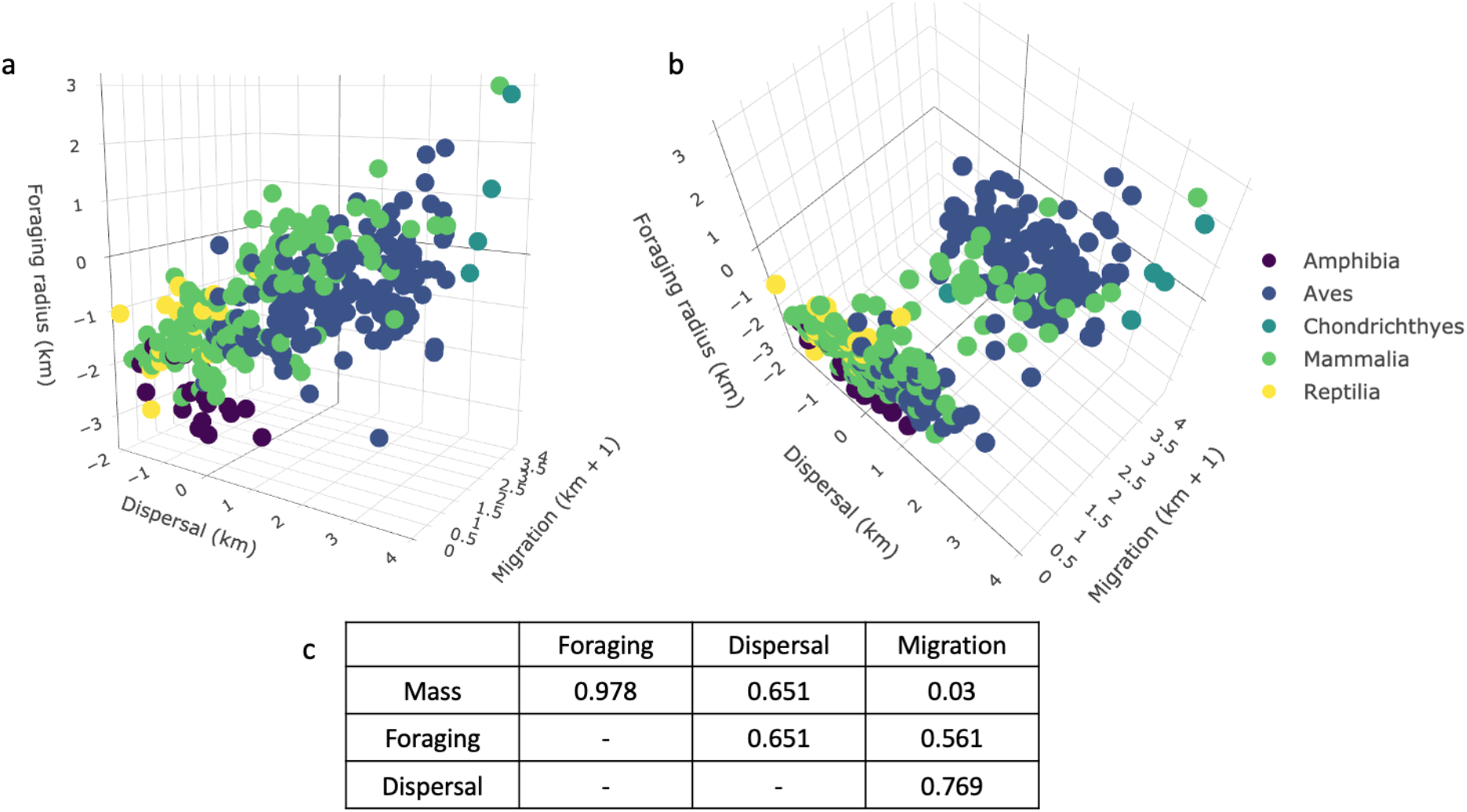
Panels (a) and (b) depict log10-transformed movement types on 3-axes for 322 species. Migration values log10+1 transformed to account for non-migratory vertebrates. Panel (c) is the covariance matrix for each movement type and body mass. See https://sdstraus.shinyapps.io/3dplot/ for interactive plot.

### Hierarchical models

Finally, we used hierarchical models to test how each hypothesized constraint modified the relationship between body size and movement distance. We found that variation in foraging and dispersal distances was partially explained by body size (M0: slopes: 0.41 ± 0.06 and 0.34 ± 0.06, respectively), while migration movement was not (slope: -0.08 ± 0.08, LOOIC: 6726.8, Figure 5, Table 2). When considering mass as a fixed effect and medium as a random effect (Models M1.1 - M1.3), we saw generally positive relationships between mass and movement for all media types for foraging and dispersal. While animals that migrate through air have no significant relationship between size and migration distance (slope: -0.09 ± 0.1, Figure 5c, Table 2), they move over larger distances than those that move in water or on land for a given body size (Figure 5a-c, Table 2).

**Figure 4.**
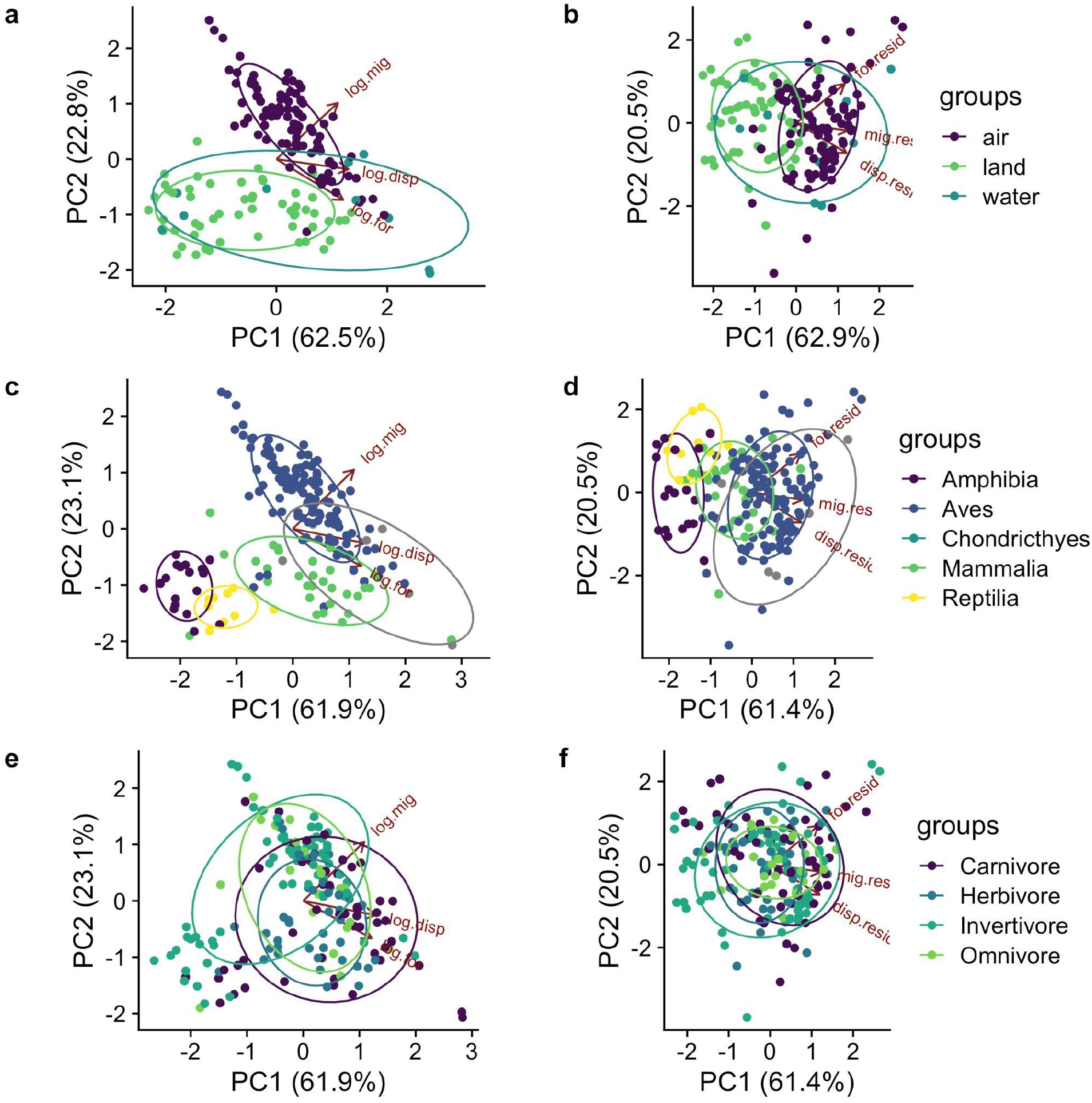
PCA plots: left column: log10 transformed movement (“log.for”, “log.disp”, “log.mig”), right column, residuals of each log10 transformed movement type a function of log10 body mass (“for.resid”, “disp.resid”, “mig.resid”). Only animals with migration greater than zero were included.

**Figure 5.**
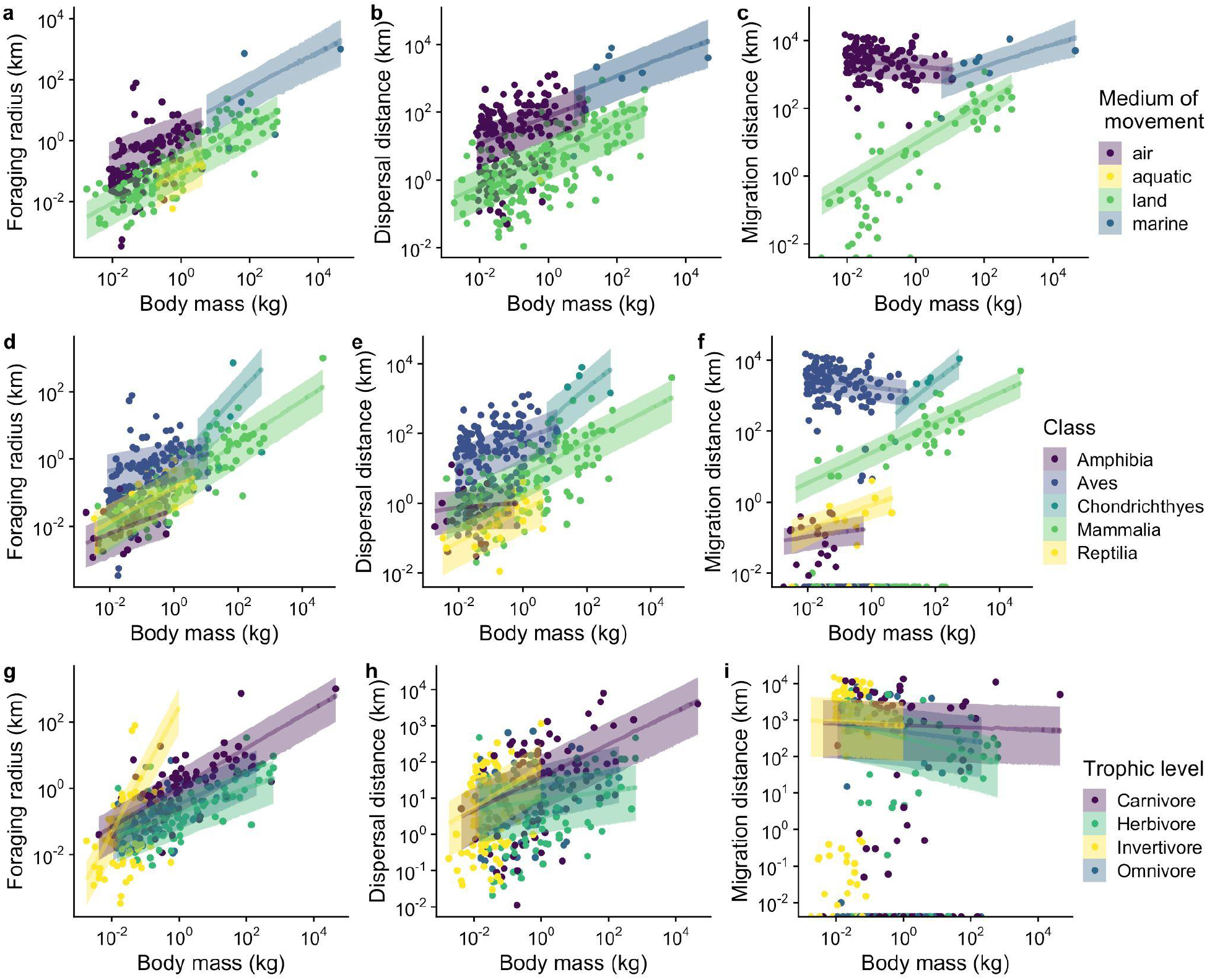
(a-c) the relationships between mass and (a) foraging, (b) dispersal, and (c) migration for three types of media: air, land and water. (d-f) the relationships between mass and (d) foraging, (e) dispersal, and (f) migration for each taxonomic class. Relationships are generally positive, except for Aves. (g-i) the relationships between mass and (g) foraging, (h) dispersal, and (i) migration for different diet categories. Relationships are positive for all diet types for foraging and dispersal, but weakly negative for migration. Shaded bands represent 50% posterior predictive probability.

Model M2, testing the shared taxonomy hypothesis, was the best performing model, with strong evidence of taxonomic constraints on movement distance (M2 LOOIC: 6068.8, Figure 5d-f, Table 2). We found that movement distance increased with body size for all taxonomic classes, with small differences in intercepts (distance moved) for each class (Table 2). However, while birds migrate farther than other classes (intercept: 7.87 ± 0.33) for a given size, we found no significant relationship between body mass and migration distance (slope: -0.11 ± 0.11, Figure 5f, Table 2). Organisms belonging to different taxonomic orders within the same class tended to move similar distances with similar body-size scaling relationships (Supplemental material, Figure S1). The order Galliformes was however one major exception to this observation (Figure S1, top right).

We found positive relationships between body size and foraging and dispersal distances for all trophic guilds except dispersing omnivores (M3 LOOIC: 6558.4, Figure 5g-h, Table 2). However, there was no relationship between body size and migration distance for most trophic guilds, with the exception that smaller herbivores migrated longer distances (slope: -0.23 ± 0.18) (Figure 5i, Table 2). While all trophic guilds dispersed and migrated similar distances (see intercepts in Table 2), we found that carnivores and invertivores had higher intercepts than other groups (1.06 ± 0.32 and 6.66 ± 1.34, respectively; Table 2).

## Discussion

Animal movement profiles structure ecological communities, but the constraints on movement, and how those constraints influence donor and receiving communities, is unknown. Here we collated a large database in order to address three possible constraints on animal movement, in addition to body size, focusing on three types of movement (foraging, dispersal and migration). Knowing the constraints on animal movement can help us to parameterize ecological models and better understand the role of movement at local, regional, and landscape scales.

We found positive covariance between the movement distances of foraging, migration and dispersal, and a strong signal of positive covariance between body size and foraging distance, moderate covariance between body size and dispersal, and weak covariance between body size and migration (Figure 3c, Table 2). Yet we also found support for our hypotheses that these relationships could be decoupled by biophysical, evolutionary and ecological constraints. Body size typically scaled positively with movement distance for all three types of movement, however the strength, slope, and even direction of that relationship was altered by each constraint considered; this was most apparent for shared taxonomy and movement media constraints (Table 2, Figure 5). However, body mass scaling of migration was much weaker within birds than other groups, where long migrations can be undertaken by small bodied animals with smaller foraging and dispersal distances.

The distribution of movement profiles that we observed could hint at constraints on movement distance. To consider this, we apply the concept of a ‘constraint envelope’ (e.g., Diniz-Filho 2004). A constraint envelope is the boundary in phenotypic space between phenotypes with values that are evolutionarily accessible or inaccessible, with ‘inaccessibilities’ arising via trade offs among functional trait dimensions and biophysical limits on any one dimension. Boundaries are difficult to ascertain with available data, such as when sampling of taxa is inexhaustive, but constraint envelopes can be interpreted in the realm of probability rather than in the realm of possibility (e.g., some phenotypes are more represented than others). Exceptional taxa (i.e., outliers that move extremely long distances compared to similarly sized species) were most strongly observed for foraging distances only, showing that it is possible to break beyond those constraints. Importantly, the constraint envelopes we observe appear to differ among taxonomic groups, leading to lumpy distributions when data are aggregated across taxa. Within taxonomic groups represented in our dataset, either biophysical limits or a lack of ecological necessity has prevented these groups from expanding their current range of movement distances, despite these expanded ranges being present in other taxa. Although we cannot know how the exact coverage of data in this trivariate volume would vary if data from more taxonomic groups were available, we nonetheless described patterns observed for the specific groups we have examined, which represents the most comprehensive dataset of movement distances to date.

### Movement Medium Hypothesis

We found that factors such as evolutionary history and the medium for locomotion modified the relationship between body size and movement. For example, small animals migrate longer distances in air compared to on land. This is evidenced by a large aggregation of points in bivariate movement space, where species have small bodies but migrate long distances, comprised of flying birds of all trophic levels (Figures 5c,f,i). Differences in viscosity, and density of water and air affect the energy used for movement (Biewener & Patek 2018). Because migration can produce a pulse of resources for receiving food webs, this influence of media on migration distance with body size may have interesting implications for meta-food webs and meta-ecosystems. For example, as the body size of animals increases, animals that migrate by air will be part of food webs that are connected at smaller spatial scales, while animals that migrate by land will connect more distant food webs. This also implies that there may be considerable variation in migration within food webs, as advocated by previous work from this research group (Guzman *et al*. 2019). The number of individuals in a migratory population will determine whether the numerical or resource subsidies provided by these species are the same: for example, fewer, larger, terrestrial migrators could connect nearby food webs and ecosystems in a similar way as many, smaller, flying migrators could connect faraway ones.

### Shared Taxonomy Hypothesis

Our analysis provides strong support for our hypothesis that evolutionary history and taxonomic constraints may limit movement distances. Shared taxonomy modified the relationship between movement distances and body size, and it accounted for more variation than either movement media or trophic guild. Evidence of the decoupling of movement types was the strongest within migration where birds tended to migrate the greatest distances. Most mammals and reptiles did not migrate, but their distribution was highly right-skewed, such that a small handful of mammalian species (primarily ungulates) migrated extremely long distances. Dispersal distance followed a similar but less extreme pattern. Distributions were relatively more normal for each group, where sharks and birds disperse the farthest, followed by mammals, reptiles and amphibians. Grouping by class is least apparent for foraging where the distribution of foraging distances are more normally distributed within and between groups. This may suggest that migration is more reflective of upper physiological limits that are evolutionarily constrained while dispersal and foraging are more labile. Dispersal and foraging could instead be constrained by, for example, population dynamics or selection on other traits (Burgess *et al*. 2016) and ecologically determined optimal foraging (Pulliam 1974), respectively, and so don’t display as strong of evolutionary signals. Supporting this idea is the total distance moved in each category, which is an order of magnitude larger for migration (Figures 2 and 3). Also supporting this assertion is the plethora of specialized adaptations in the longest distance migrators (Weber 2009), though these adaptations are fairly labile within clades of birds (Pulido 2007). For example, in our dataset, the bird Order Galliformes, had a significantly lower migration intercept than other groups (Figure S1). This group, containing pheasants, prairie chickens, and other land fowl, are characterized by high wing-loading and low wing aspect ratios (Rayner 1988), making them relatively weak flyers.

### Trophic Guild Hypothesis

We found that trophic guild modified covariance between movement and body size to a lesser extent than either media or shared taxonomy, but the model still performed better than the null according to our model selection (Table 2). Our analysis found that both dispersal and foraging distances increased with body size for all guilds except for omnivorous dispersers (Table 2, Figure 5g-i). However, body sizes are not evenly distributed among the trophic guilds. In our dataset, invertivores had the smallest body sizes, omnivores and herbivores slightly larger, and carnivores achieving the largest body sizes. This is contrast somewhat with the findings of Potapov et al. (2019) who found a relationship between trophic level and body size for marine organisms, but not terrestrial or freshwater. However, the Potapov et al. (2019) study also considers invertebrates, including zoo and phytoplanktons, while our study only considers vertebrates. Invertivores had a significantly steeper relationship between body size foraging movement than all other groups, and carnivores had a steeper slope than herbivores, but not omnivores. The substantially steeper slope and higher intercept observed in invertivores compared to other trophic guilds was driven primarily by *Apus apus* (Common swift) and *Progne subis* (Purple martin), two small-bodied birds that forage over long distances (Table 2, Figure 3, Figure 5g). For dispersal (Figure 5h), invertivores and carnivores had approximately similar slopes, both of which were significantly steeper than those for herbivores and omnivores. Conversely, all slopes were negative for migration (Figure 5i), such that smaller organisms migrate farther than larger ones, but this relationship was only significant for herbivores. Unlike our other two analyses described above, where aerial movement occurs primarily by birds (Aves), groupings are influenced by shared evolutionary history. This shared history is either explicit (i.e. class as grouping factor) or implicit (i.e. media as grouping factor, where more closely related species are more likely to use similar media). Trophic guild, while not entirely independent of shared evolutionary history, is less influenced by it. As such, we don’t see distinct groupings of points for trophic guild as we do for media or taxonomy (Figure 3 interactive plot, Figure 4d). In our PCA, those groupings become even less distinct when we account for body mass (Figure 4f), whereas there is substantially less overlap in groups when considering either media or shared evolutionary history.

### Exceptions, caveats, and implications

Finally, we can further understand constraints by examining exceptional species in our dataset, which reveal the potential importance of factors we did not consider or measure in our analysis. For some species, exceptions to general observations may be bounded by physical limits. For example, sperm whales have enormous foraging ranges, spanning 2000 kilometers, whereas other, smaller species (e.g., migratory birds) migrate on average ∼77,000x farther than the diameter of their foraging range (a staggering ratio). If a sperm whale migrated at distances this many times greater than its foraging range, migration would exceed the circumference of the Earth. For some other species, exceptions may be driven by behavior. Some birds (as well as non-avian animals) have colonial (or social) lifestyles or exhibit philopatry, thus reducing their necessary dispersal distances through behavioral adaptations. Lastly, some deviations from general relationships may also result from underlying environmental conditions (e.g., NDVI Pettorelli *et al*. 2011). For example, two animals of the same body size may need larger or smaller foraging ranges depending on the productivity of their environments. However, this environmental constraint on food availability in their foraging ranges may not affect the distance they disperse or migrate.

While our study represents the largest assembled database on animal movement, to date, our results are still limited by the fact that not all taxa could be included. We were able to find dispersal, migration or foraging distances for many more species, but the number of species that had all three forms of movement characterized was limited. Notably, the data set we compiled does not contain any invertebrates or passive dispersers. These two types of organisms would likely drastically change the covariance relationships we observed here. Using mean values of movement may also hide interesting variation within species, e.g. facultative versus obligate migration (Newton 2012). Additionally, we excluded nomadic animals from this dataset because nomadism is a movement strategy that is understudied while also combining characteristics of the other modes of movement. Nomadism, where animals make seemingly random, long distance movements, typically occurs in resource variable environments and should also influence ecological dynamics (Teitelbaum & Mueller 2019). Finally, movement may also be constrained by traits that we did not consider in this study. For example, recent work by Esmaeili et al. (2021) found that digestive strategy in large mammals, i.e., fore-versus hind-gut fermentation, influences patterns of landscape use. We hope the present study will encourage researchers to measure these forms of movement for other types of organisms, and to expand on the types of movement considered.

Describing species’ movement profiles might help advance and simplify an ecological understanding that integrates across scales. Because movement types differ in how they manifest through both space and time, each is expected to have unique contributions to ecological dynamics (Box 1). An exploration of the dynamics that might emerge depending on movement profiles of interacting species in communities is a fruitful avenue of future research, and modeling efforts could benefit from a rough baseline to parameterize movement rates and distances in models, as we provide here. We show that there are no universal scaling relationships between movement types — for individual taxa, movement distances depend on movement type, in ways that reflect body size, evolutionary history, trophic level, and movement media. However, we do show boundaries on movement distance, helping to narrow in estimates, as well as differences in overall order of magnitude between movement types. We expect empirical estimates to be particularly useful in extending spatial diversity theories (e.g., metacommunity ecology, spatial coexistence theory) into a multi-trophic context, where interactions (e.g., predation, mutualism) are often realized between species with vastly different body sizes, ecologies, and evolutionary histories. These theories have made monumental contributions to our understanding of spatial processes and local-regional coupled dynamics, but until recently have left aside information about foraging and migration (Guzman *et al*. 2019).

### Conclusion

In sum, we explored how dispersal combines with migration and foraging movements and body size to create species’ movement profiles. The distribution of movement profiles across taxa in phenotypic trait space interact with physiological, taxonomic, and ecological traits to create constraint envelopes that signal the boundary of what movement distances may be possible. Our study not only provides insights into the parameterization of future ecological models, but also captures a holistic view of animal movement that is missed when considering each movement type in isolation.

As movement ecologists seek greater integration with conservation (Fraser *et al*. 2018), understanding how the myriad types of movement can better inform management and restoration efforts will be key. The generalized patterns we describe here can thus augment movement information on data poor species. This information is needed for the design of protected areas (Noonan *et al*. 2020), wildlife corridors (Ford *et al*. 2020), the spread of pathogens (Nobert *et al*. 2016), and new formulations of critical habitat designations (Davy *et al*. 2017).

## Acknowledgements

Working group funding was provided by UBC through a Research Excellence Cluster grant for Catalyzing Biodiversity Research. S.S. was supported by NSERC Discovery Grants and UBC One Year Fellowships. P.L.T and C.J.L. were supported by NSERC and Killam postdoctoral fellowships. M.I.O’C, D.G. A.T.F, C.J.L. and R.M.G. are supported by NSERC Discovery Grants, and L.M.G. is supported by the Liber Ero Fellowship. The authors thank Dr. Matthew Pennell for his input on how to analyze the effect of shared phylogeny. The authors have no conflicts of interest to declare.

## Authorship statement

Data were collected by SS, LMG, DAM, CJL, CF, with support from PLT and RG. LMG and SS analyzed the data, with support from PLT. First draft of the manuscript was written by SS, LMG, CJL, CF, RG, and DAM, with support from MO. All authors participated in discussions and subsequent revisions of the manuscript.

