## Supplemental material for "Macroecological variation in movement profiles: body size does not explain it all"

**This file includes**

**Supplementary Figures**

**Figure S1.** Relationships between body size and movement for each taxonomic order

**Supplementary Tables**

**Table S1. PCA loadings for movement types**

**Supplementary Appendices**

**Appendix S1.** Full Model Formulations

**Appendix S2.** Data synthesis literature cited

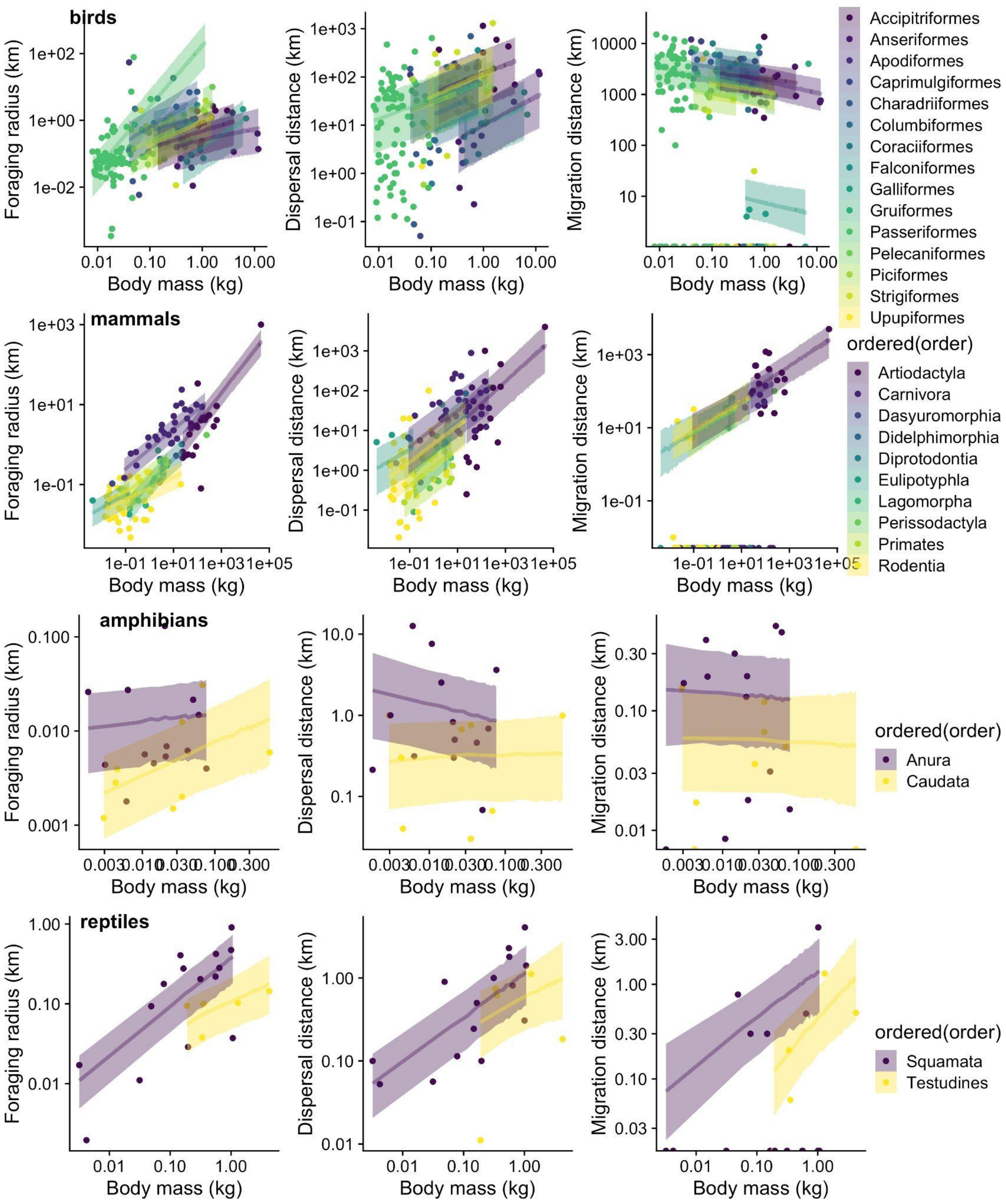
birds
mammals
amphibians
reptiles
Foraging radius (km)
Dispersal distance (km)
Migration distance (km)
Body mass (kg)
Accipitriformes
Anseriformes
Apodiformes
Caprimulgiformes
Charadriiformes
Columbiformes
Coraciiformes
Falconiformes
Galliformes
Gruiformes
Passeriformes
Pelecaniformes
Piciformes
Strigiformes
Upupiformes
ordered(order)
Artiodactyla
Carnivora
Dasyuromorphia
Didelphimorphia
Diprotodontia
Eulipotyphla
Lagomorpha
Perissodactyla
Primates
Rodentia
ordered(order)
Anura
Caudata
ordered(order)
Squamata
Testudines

**Figure S1.** The relationships between mass and foraging, dispersal, and migration for taxonomic order within class. For birds, the relationship between body mass and foraging and dispersal distances were generally weakly positive for all orders, while weakly negative for migration. The order Galliformes had migration distances two orders of magnitude less than most other orders. For mammals, movement distances were tightly linked to body size for all types of movement. Within amphibians, correlations we found no relationship between body size and movement distance for any order or any movement type, while we found weak relationships within orders for reptiles. Shaded bands represent 50% posterior predictive probability.

**Table S1.** PCA loadings for log10 transformed movement, and the residuals of log10 transformed movement as a function of body mass.

| **Movement (Log10)** | **PC1** | **PC2** |
|---|---|---|
| Foraging | 0.58 | -0.52 |
| Dispersal | 0.62 | -0.20 |
| Migration | 0.52 | 0.83 |
| residuals(Foraging) | 0.56 | 0.77 |
| residuals(Dispersal) | 0.56 | -0.61 |
| residuals(Migration | 0.60 | -0.15 |

**Supplementary Appendices**

**Appendix S1:** Full hierarchical model configurations

Model 0 tests the effects of body mass. The same error structure was also used for models 1-3:

$$dispersal_i = \beta_0 + \beta_{body\ size[i]} + \varepsilon d_i$$
$$\varepsilon d_i \sim Gamma(\mu, \kappa)$$

$$foraging_i = \beta_0 + \beta_{body\ size[i]} + \varepsilon h_i$$
$$\varepsilon h_i \sim Gamma(\mu, \kappa)$$

$$migration_i = \beta_0 + \beta_{body\ size[i]} + \varepsilon m_i$$
$$\varepsilon m_i \sim HurdleGamma(\mu, \kappa, \eta)$$

$M0 = M[dispersal_i, foraging_i, migration_i]$

Models 1 tests the effects of media:

$$foraging_i = \beta_0 + \beta_{body\ size} + \alpha_{0media} + \alpha_{body\ size\ media}$$
$$dispersal_i = \beta_0 + \beta_{body\ size} + \alpha_{0media} + \alpha_{body\ size\ media}$$
$$migration_i = \beta_0 + \beta_{body\ size} + \alpha_{0media} + \alpha_{body\ size\ media}$$

$M1.1 = dispersal_i, M1.2 = foraging_i, M1.3 = migration_i$

Models 2 tests the effects of taxonomy through class (2.1) or order (2.2). For the main figures and tables we present the results of Model 2.1 (class), and we present results for model 2.2 in the supplementary materials.

Model 2.1

$$foraging_i = \beta_0 + \beta_{body\ size} + \alpha_{0class} + \alpha_{body\ size\ class}$$
$$dispersal_i = \beta_0 + \beta_{body\ size} + \alpha_{0class} + \alpha_{body\ size\ class}$$
$$migration_i = \beta_0 + \beta_{body\ size} + \alpha_{0class} + \alpha_{body\ size\ class}$$

$M2.1 = M[dispersal_i, foraging_i, migration_i]$

Model 2.2

$$foraging_i = \beta_0 + \beta_{body\ size} + \alpha_{0order} + \alpha_{body\ size\ order}$$
$$dispersal_i = \beta_0 + \beta_{body\ size} + \alpha_{0order} + \alpha_{body\ size\ order}$$
$$migration_i = \beta_0 + \beta_{body\ size} + \alpha_{0order} + \alpha_{body\ size\ order}$$
$$M2.2 = M[dispersal_i, foraging_i, migration_i]$$

Model 3 tests the effects of trophic guild:

$$foraging_i = \beta_0 + \beta_{body\ size} + \alpha_{0guild} + \alpha_{body\ size\ guild}$$
$$dispersal_i = \beta_0 + \beta_{body\ size} + \alpha_{0guild} + \alpha_{body\ size\ guild}$$
$$migration_i = \beta_0 + \beta_{body\ size} + \alpha_{0guild} + \alpha_{body\ size\ guild}$$
$$M3 = M[dispersal_i, foraging_i, migration_i]$$

**Appendix 2:** Data synthesis literature cited

### Dispersal

11. Brigham, R. M., J.Ng, R. G. Poulin, and S. D.Grindal. 2011. Common Nighthawk (Chordeiles minor), version 2.0. in A. F. Poole, ed. In the birds of North America, Cornell Lab of Ornithology, Ithaca, New York, USA.

23. Georgiadis, N. (1995). Population structure of wildebeest: implications for conservation. *Serengeti II: Dynamics, Management, and Conservation of an Ecosystem*, *2*, 473.

24. Giovanni, M. D. (2009). *Demographics and habitat selection for the western meadowlark (Sturnella neglecta) in the Nebraska sandhills* (L. A. Powell & W. H. Schacht (eds.)) [The University of Nebraska - Lincoln].

25. Hayes, M. A. (2015). *Dispersal and population genetic structure in two flyways of sandhill cranes (Grus canadensis)* (M. E. Berres & J. A. Barzen (eds.)) [The University of Wisconsin - Madison].

26. Hofer, H., & East, M. (1995). Population dynamics, population size, and the commuting system of Serengeti spotted hyenas. *Serengeti II: Dynamics, Management, and Conservation of an Ecosystem*, *2*, 332.

32. Kus, B., Hopp, S. L., Johnson, R. R., Brown, B. T., & Poole, A. F. (2020). Bell's Vireo (Vireo bellii), version 1.0. *Birds of the World (AF Poole, Editor). Cornell Lab of Ornithology, Ithaca, NY, USA.*

61. van Oort Bruce N. McLellan and Robert Serrouya, H. (2008). *Dispersal in a declining caribou "meta-population"?*

https://www.env.gov.bc.ca/wildlife/wsi/reports/4442_WSI_4442_RPT_DISPERSALPAPER.DOC

62. Vines, T. H. (2002). *Migration, habitat choice and assortative mating in a Bombina hybrid zone* [zoology.ubc.ca]. http://www.zoology.ubc.ca/~vines/r/pdf/VinesThesis.pdf

63. Watson, J. W., Banasch, U., Byer, T., Svingen, D. N., McCready, R., Hanni, D., & Gerhardt, R. (2019). First-Year Migration and Natal Region Fidelity of Immature Ferruginous Hawks. *The Journal of Raptor Research*, *53*(3), 266–275. https://doi.org/10.3356/JRR-18-32

64. Whitmee, S., & Orme, C. D. L. (2013). Predicting dispersal distance in mammals: a trait-based approach. *The Journal of Animal Ecology*, *82*(1), 211–221. https://doi.org/10.1111/j.1365-2656.2012.02030.x

### Home Range

18. Cygnus olor. (n.d.). Global Invasive Species Database. http://www.iucngisd.org/gisd/speciesname/Cygnus+olor

35. Guangmei, P. C. Z. (2003). A study on home range of tree sparrow (Passer montanus) in Beijing Normal University in winter. Journal of Beijing Normal University (Natural Science).

41. Helon, D. A. (2006). Summer home range, habitat use, movements, and activity patterns of river otters (Lontra canadensis) in the Killbuck Watershed, northeastern Ohio. West Virginia University.

42. Heupel, M. R., Simpfendorfer, C. A., & Hueter, R. E. (2004). Estimation of Shark Home Ranges using Passive Monitoring Techniques. *Environmental Biology of Fishes, 71*(2), 135–142. https://doi.org/10.1023/B:EBFI.0000045710.18997.f7

43. Hillis, R. E., & Bellis, E. D. (1971). Some Aspects of the Ecology of the Hellbender, Cryptobranchus alleganiensis alleganiensis, in a Pennsylvania Stream. *Journal of Herpetology, 5*(3/4), 121–126. https://doi.org/10.2307/1562734

44. Holmes, R. T. (1973). Social behaviour of breeding western sandpipers Calidris mauri. *The Ibis, 115*(1), 107–123. https://doi.org/10.1111/j.1474-919X.1973.tb02627.x

45. Hooper, R. G., Niles, L. J., Harlow, R. F., & Wood, G. W. (1982). Home Ranges of Red-Cockaded Woodpeckers in Coastal South Carolina. *The Auk, 99*(4), 675–682. https://doi.org/10.1093/auk/99.4.675

46. Ivey, G. L., Dugger, B. D., Casazza, M. L., Fleskes, J. P., & Herziger, C. P. (2011). Movements and Home Range Size of Greater and Lesser Sandhill Cranes Wintering in Central California. Twelfth North American Crane Workshop, 94.

67. Martin, M. (2016). Winter Ecology and Behavior of Eastern Towhees at Taylor Fork Ecological Area.

84. Renfrew, Rosalind, Allan M. Strong, Noah G. Perlut, Stephen G. Martin andThomas A. Gavin. (2015). Bobolink (Dolichonyx oryzivorus), The Birds of North America (P. G. Rodewald, Ed.).

Ithaca: Cornell Lab of Ornithology; Retrieved from the Birds of North America: https://birdsna.org/SpeciesAccount/bna/species/boboli DOI: 10.2173/bna.176

ifornia_Wildlife_Habitat_Relationships_System_California_Department_of_Fish_and_Wildlife/links/587fd7f108ae9275d4ee3a7c/Spotted-Sandpiper-Actitis-macularius-Species-Account-in-California-Wildlife-Habitat-Relationships-System-California-Department-of-Fish-and-Wildlife.pdf

**Body Mass**

### Migration

1. Ahlborn, G & White, M. (1999). American Marten. California Department of Fish and Game, California's Wildlife, Sacramento, CA. http://www.sibr.com/mammals/M154.html. Accessed September 10, 2021

2. Ahlborn, G., & White, M. Fisher. (2005). Martes pennanti  California Department of Fish and Game. California Interagency Wildlife Task Group. California Wildlife Habitat Relationships version 8.1 personal computer program. Sacramento, California.

11. Baskett, T. S., Sayre, M. W., & Tomlinson, R. E. (Eds.). (1993). Ecology and management of the mourning dove. Stackpole Books.

12. Beauvais, G. P., & Dark-Smiley, D. N. (2005). Species Assessment for Idaho Pocket Gopher (Thomomys idahoensis) in Wyoming. *Unpublished Report. Wyoming Natural Diversity Database, University of Wyoming, Laramie.* http://www.uwyo.edu/wyndd/_files/docs/reports/speciesassessments/idahopocketgopher-jun2005.pdf

    Beauvais, G. P., & Johnson, L. (2004). Species assessment for wolverine (Gulo gulo) in Wyoming. *US Department of Interior, Bureau of Land Management, Cheyenne, Wyoming,* 20.

58. Eccles, K. M., Thomas, P. J., & Chan, H. M. (2020). Relationships between mercury concentrations in fur and stomach contents of river otter (Lontra canadensis) and mink (Neovison

vison) in Northern Alberta Canada and their applications as proxies for environmental factors determining mercury bioavailability. *Environmental Research, 181*, 108961. https://doi.org/10.1016/j.envres.2019.108961

68. Gonzales, A. G. (1997). Weasels, skunks, ringtail, raccoon, and badger. *The Wildlife Society California North Coast Chapter*, 63.

84. Horton, K. G., Van Doren, B. M., La Sorte, F. A., Fink, D., Sheldon, D., Farnsworth, A., & Kelly, J. F. (2018). Navigating north: how body mass and winds shape avian flight behaviours

across a North American migratory flyway. *Ecology Letters, 21*(7), 1055–1064. https://doi.org/10.1111/ele.12971

85. Howell, A. H., Jackson, H. H. T., Murie, O. J., & Bailey, V. (1931). Revision of the American Chipmunks:(genera Tamias and Eutamias) (No. 51-54). US Government Printing Office.

86. Huntly, N. J., Smith, A. T., & Ivins, B. L. (1986). Foraging behavior of the pika (Ochotona princeps), with comparisons of grazing versus haying. *Journal of Mammalogy*, 67(1), 139-148.

87. Jacobs, J. P., & Jacobs, E. A. (2002). Conservation assessment for red-shouldered hawk (Buteo lineatus): National Forests of north central states. US Department of Agriculture, Forest Service, Eastern Region, Milwaukee, Wisconsin.

88. Johnson, J. A., Booms, T. L., DeCicco, L. H., & Douglas, D. C. (2017). Seasonal movements of the Short-eared Owl (Asio flammeus) in western North America as revealed by satellite telemetry. *Journal of Raptor Research*, 51(2), 115-128.

89. Johnston, B. (1999). Terrestrial Pacific Giant Salamanders (Dicamptodon tenebrosus Good): natural history and their response to forest practices (Doctoral dissertation, University of British Columbia).

123. Mitchell, C. D. (1994). Trumpeter swan (Cygnus buccinator). *The Birds of North America*.

136. Razafindratsima, O. H., Yacoby, Y., & Park, D. S. (2018). MADA: Malagasy animal trait data archive.

137. Reading, C., & Jofré, G. (2009). Habitat selection and range size of grass snakes Natrix natrix in an agricultural landscape in southern England. *Amphibia-Reptilia*, 30(3), 379-388.

138. Reed, J. M., Oring, L. W., Gray, E. M., & Rodewald, P. G. (2013). Spotted sandpiper (Actitis macularius). *The Birds of North America Online. Ithaca, NY: Cornell Lab of Ornithology*.

148. Sibley, D., Elphick, C., & Dunning, J. B. (2001). *The Sibley guide to bird life & behavior* (No. Sirsi) i9780679451235). National Audubon Society.

149. Sidorov, G. N., & Putin, A. V. (2010). The house mouse (Mus musculus L.) in Omsk educational institutions: Seasonal migration, abundance, reproduction, distribution, foraging, and associated damage. *Contemporary Problems of Ecology, 3*(5), 601–605. https://doi.org/10.1134/S1995425510050164

150. Smith, W. P. (2007). Ecology of Glaucomys sabrinus: Habitat, Demography, and Community Relations. *Journal of Mammalogy, 88*(4), 862–881. https://doi.org/10.1644/06-MAMM-S-371R1.1

151. Söderquist, P., Gunnarsson, G., & Elmberg, J. (2013). Longevity and migration distance differ between wild and hand-reared mallards Anas platyrhynchos in Northern Europe. European *Journal of Wildlife Research*, 59(2), 159-166.

152. Squires, J. G., & Reynolds, R. T. (1997). Northern Goshawk (Accipiter gentilis) In: Poole A, Gill F (eds) The birds of North America, No 298. The Academy of Natural Sciences, Philadelphia, and the American Ornithologists' Union, Washington DC.

158. Sunquist, M. E. (1979). The movement and activities of tigers (Panthera tigris) in Chitwan National Park, Nepal (Doctoral dissertation, Ph. D. dissertation, University of Minnesota).

159. Sunquist, M.E., & Eisenberg, J.F. (2016). Reproductive strategies of female *Didelphis.*

172. Verbeek, N. A., & Caffrey, C. (2002). *American Crow: Corvus Brachyrhynchos*. Birds of North America, Incorporated.

173. Verts, B. J., & Carraway, L. N. (2001). Tamias minimus. *Mammalian Species, 2001*(653), 1–10. https://doi.org/10.1644/1545-1410(2001)653<0001:TM>2.0.CO;2

174. Vuarin, P., Dammhahn, M., Kappeler, P. M., & Henry, P. Y. (2015). When to initiate torpor use? Food availability times the transition to winter phenotype in a tropical heterotherm. *Oecologia*, *179*(1), 43-53.

175. Warkentin, I. G., & Oliphant, L. W. (1990). Habitat use and foraging behaviour of urban merlins (Falco columbarius) in winter. *Journal of Zoology*, *221*(4), 539-563.

176. Wassmer, T., & Refinetti, R. (2016). Daily activity and nest occupation patterns of fox squirrels (Sciurus niger) throughout the year. *PloS one*, *11*(3), e0151249.

### Diet

### Media

28. Kear, J. (2005). *Ducks, Geese and Swans: Species accounts (Cairina to Mergus)*. Oxford University Press.
